# Investigating the Role and Regulation of GPNMB in Progranulin-deficient Macrophages

**DOI:** 10.1101/2024.03.12.584649

**Authors:** Drew A. Gillett, Noelle K. Neighbarger, Cassandra Cole, Rebecca L. Wallings, Malú Gámez Tansey

## Abstract

Progranulin is a holoprotein that is critical for successful aging, and insufficient levels of progranulin are associated with increased risk for developing age-related neurodegenerative diseases like AD, PD, and FTD. Symptoms can vary widely, but a uniting feature among these different neurodegenerative diseases is prodromal peripheral immune cell phenotypes. However, there remains considerable gaps in the understanding of the function(s) of progranulin in immune cells, and recent work has identified a novel target candidate called GPNMB. We addressed this gap by investigating the peritoneal macrophages of 5-6-month-old *Grn* KO mice, and we discovered that GPNMB is actively increased as a result of insufficient progranulin and that MITF, a transcription factor, is also dysregulated in progranulin-deficient macrophages. These findings highlight the importance of early-stage disease mechanism(s) in peripheral cell populations that may lead to viable treatment strategies to delay disease progression at an early, prodromal timepoint and extend therapeutic windows.

## Introduction

Progranulin is a glycosylated holoprotein that is composed of seven and a half cysteine-enriched repeats, called granulins, that are distributed as “beads on a string” with short linker regions joining the repeats together^1^. Nascent progranulin is translated into the endoplasmic reticulum (ER), where ER-resident chaperone and folding proteins aid in the cysteine-cysteine disulfide bonds that permit the beta-fold structure of the granulins. After glycosylation, progranulin can be secreted out of the cell or intracellularly trafficked to the endo-lysosomal pathway directly. Extracellular progranulin enters the cell via interaction with its canonical receptor, sortilin (Sort1)^2^, or by piggy-backing onto a similar holoprotein, prosaposin, which is internalized with LRP1 and M6PR^3^. Secreted progranulin circulates throughout the body fluids, including blood serum and cerebrospinal fluid (CSF) as a dimer^4^. In this way, the level of progranulin expression is spread throughout the body, allowing even low-producing cells to take up sufficient amounts of progranulin.

Progranulin is critical for successful aging^5^. Multiple mechanisms can influence progranulin expression, but loss of full progranulin expression, approximately 96-125ng per mL of blood serum^6,7^, can increase risk of disease. The silencing of one *GRN* allele is associated with an increased risk for Frontotemporal Dementia (FTD), but the silencing of both alleles results in Neuronal Ceroid Lipofuscinosis (NCL-11)^8,9^. Furthermore, other age-related neurodegenerative diseases, like Parkinson’s Disease (PD) and Alzheimer’s Disease (AD), are associated with decreased progranulin levels^10-18^, suggesting common underlying mechanisms regarding the role of progranulin in healthy aging.

The specific functions of progranulin are multi-faceted and impact specific cell populations differently. Peripheral immune cells rely on progranulin. Clinically-symptomatic *GRN*-mutation carriers showed increased soluble Cluster of Differentiation 163 (CD163) and CC Chemokine Ligand 18 (CCL18) serum levels relative to healthy controls, suggesting increased peripheral macrophage activity^19^. In addition, lipopolysaccharide binding protein (LBP) levels correlated with white-matter changes in the frontal lobe and Clinical Dementia Rating (CDR)-FTLD sum of boxes (SB) scores of clinically symptomatic *GRN*-FTD patients, indicating that peripheral immune activity correlates with region-specific brain changes and clinically relevant behavior. When macrophages isolated from progranulin-deficient mice were challenged with lipopolysaccharide (LPS), they showed increased transcription of *Mcp-1*, *Il-12p40*, and *Tnf,* as well as decreased *Il-10* transcription, relative to wild-type (WT) controls^20^ (Yin et al. 2009).

In the brain, progranulin is associated with neuroprotection. The precise mechanism(s) are unclear, but progranulin modulates neuroinflammation and reduces brain volume loss in stroke and head injury paradigms^21,22^ and also prevented dopaminergic neuron loss in an MPTP model^23^. Recent work identified a specific increase in the expression of glycoprotein non-metastatic B (GPNMB) in *GRN*-FTD patient brain lysate relative to both non-demented controls and other FTD related mutations^24^. In addition, CSF from *GRN*-FTD patients showed increased amounts of GPNMB relative to non-demented controls^24^. This indicates a specific increase in GPNMB expression with the loss of progranulin. When a mouse model of progranulin deficiency was investigated, an age-dependent increase in the expression of GPNMB was discovered. Brain lysate from young 3-month-old *Grn* knock-out (KO) mice did not show a significant change relative to age-matched wild-type (WT) controls, but 18-month old *Grn* KO mice replicated the increase in GPNMB expression seen in the *GRN*-FTD patients^24^. This gives further support that extended periods of insufficient progranulin results in increased GPNMB expression, likely in a compensatory mechanism(s) related to endo-lysosomal dysfunction.

Originally identified in a cancer cell line^25^, GPNMB is a glycoprotein that is present both on the cell membrane and intracellularly in the autophagy pathway, and is highly expressed on myeloid cells. ADAM/MMPs can cleave off the extracellular domain, generating a soluble extracellular fragment (ECF)^26^, which has been demonstrated to serve a signaling role in a paracrine and an autocrine manner^27,28^. CD44 and Syndecan-4 are the canonical cell-surface receptors for GPNMB, but the effect of binding the GPNMB ECF is cell-type dependent^29,30^. Within the cell, GPNMB localizes with LC3+ vesicles, where it is believed to support the recruitment of the lysosome to autophagosomes^31^, allowing for autophagic cargo to be successfully degraded. Given that progranulin has a profound effect on the endo-lysosomal pathway, it seems logically consistent that GPNMB would increase in an attempt to aid autophagolysosomal function. However, the mechanisms behind GPNMB regulation are not entirely well known. Microphthalmia-associated transcription factor (MITF) has been demonstrated to regulate GPNMB in dendritic cells^32^, a class of immune cells with similar properties to macrophages.

Under basal conditions, mTORC1 phosphorylates the MiT/TFE transcription factor family of proteins (MITF, TFE3, TFEB, and TFEC), preventing their translocation to the nucleus by promoting interaction with the 14-3-3 cytosolic protein^33-35^. In conditions with decreased mTORC1 activity, such as serum starvation and/or endo-lysosomal dysfunction, MiT/TFE transcription factors are not phosphorylated by mTORC1 and can drive the expression of CLEAR (Coordinated Lysosomal Expression and Regulation) genes^36^, that push the cell towards increased autophagy and endo-lysosomal activity in an attempt to regain cellular homeostasis. However, there are a number of unknowns surrounding if and how these elements interact to contribute to the age-dependent increase in GPNMB expression in myeloid cells. This work seeks to close these gaps in knowledge by extending the investigation of GPNMB expression into the peripheral immune system and exploring candidate regulatory mechanism(s) that have been implicated to regulate GPNMB expression in the context of progranulin-deficiency.

## Materials and Methods

### Animals

C57BL/B6J mice and *Grn* KO^20^ (The Jackson Laboratory, strain #:013175) mice were generated and maintained in the McKnight Brain Institute vivarium (University of Florida) at 22°C at 60-70% humidity with a 12-hour light/dark cycle and *ad libitum* access to standard rodent chow and water. C57BL/6J (B6) (Strain #000664) controls were used for all studies. All animal procedures were approved by the University of Florida Institutional Animal Care and Use Committee and were in accordance with the National Institute of Health Guide for the Care and Use of Laboratory Animals (NIH Publications No. 80-23, 8^th^ edition, revised 2011). Mice of both genotypes were co-housed and aged to 5-6 months.

On Day 1, WT and *Grn* KO mice were injected with 1mL of sterile 3% (m/v) BBL Thioglycollate Medium Brewer Modified (Becton Dickson, 211716) in diH2O. Mice also received Buprenorphine Sustained-Release (Zoopharm; Windsor, CO) every 48 hours for pain relief. On day 4, mice were sacrificed by cervical dislocation and their peritoneal membrane was exposed by carefully dissecting the skin and fur away from the midline. Great care was taken to avoid disrupting the integrity of the peritoneal membrane. 10mL of ice-cold RPMI media (Gibco #11875) was injected into the peritoneal cavity with a 27g syringe and gentle shaking was applied to loosen adherent cells into suspension. The media was then aspirated out with a 25g syringe and dispensed into a 15mL conical and placed on ice. After all of the cells were collected, the cell suspensions were filtered through a 70um pre-wet filter overlaid a 50mL conical and rinsed twice with ice-cold 1x HBSS -/- (Gibco #14175). The resulting suspension was spun at 400 x *g* for 5 minutes at 4°C. To remove red blood cells that may have contaminated the peritoneal cell suspension, the cell pellet was treated with 2mL of ACK Lysis Buffer (0.15 M NH_4_Cl 10mM KHCO_3_ 0.1mM EDTA) and gently vortexed until the pellet was resuspended. The ACK buffer was allowed to incubate until the red blood cells lysed, approximately 1-2 minutes. The reaction was quenched with 5mL of HBSS -/-, and the cell suspension was spun again at 400 x *g* for 5 minutes at 4°C. The resulting pellets were transferred into a sterile hood and the HBSS-ACK supernatant was aspirated away. The cells were resuspended in 3mL of warm complete RPMI media (Gibco RPMI #14175, 10% FBS, 1x Pen/Strep (Gibco #15140)) and counted and viability recorded using trypan-blue exclusion on an automated cell-counter (Countess™; Thermo). 2×10^6^ cells were plated in each well of a 6-well dish. Cells were incubated at 37°C, 5% CO_2_ for five to six hours to allow macrophages to adhere. Wells were washed twice with warm 1x DPBS -/- (Gibco #14190) to remove non-adherent peritoneal cells, leaving the adherent peritoneal macrophages behind. Warm complete RPMI media was applied to the macrophages until the experimental paradigm began the following day.

### pMac Treatment Paradigm

Recombinant mouse GPNMB ECD (Aviscera Biosciences, product code 00719-03-10) was diluted in pre-warmed complete RPMI to a final concentration of 0.5ug/ml and 1.0ug/mL and was applied for 4 hours prior to lipopolysaccharide (LPS, from *Escherichia coli* O111:B4, Sigma-Aldrich) treatment. LPS-only and no-treatment controls were given a media change of fresh media at the same time. LPS was diluted to a final concentration of 1ug/mL, approximately 3,000 endotoxin units (EU), in pre-warmed complete RPMI media and applied to the macrophages for 3 hours. No treatment controls received an additional media change at the same time. Recombinant mouse progranulin (R&D Systems, catalog #: 2557-PG) was diluted in complete RPMI media to a final concentration of 5 ug/mL and applied for 48 hours. No treatment controls received a media change at the same time.

### Protein Fractionation and Western Blotting

At the end of the experimental paradigm(s), the plates containing the macrophages were removed from the incubator and placed on ice. Media was collected from select groups for downstream analysis before the macrophages were rinsed twice with ice-cold 1x PBS. After the PBS was aspirated off the second time, the cells were scraped up into the cytoplasmic fractionation buffer (150 mM sodium chloride, 50 mM Tris, pH 8.0, and 0.5% Triton X-100) with 1x protease and phosphatase inhibitors (c0mplete™ Protease Inhibitor Cocktail, Roche, product number 04693159001; PhosSTOP™, Roche, product number 04906837001). The resulting lysate was transferred to a labeled 1.5mL tube and placed on ice. Cell lysate was spun at 20,000 x *g* for 20 minutes at 4°C. After the spin was complete, the supernatant containing the soluble cytoplasmic proteins was transferred to a new labeled 1.5mL tube. The insoluble pellet was washed by adding the same volume of cytoplasmic fractionation buffer and vortexed briefly before being spun again at 13,000 x *g* for 20 minutes at 4°C. The supernatant was aspirated off, and the nuclear fractionation buffer (150 mM sodium chloride, 1.0% NP-40, 0.5% sodium deoxycholate, 0.1% SDS, and 50 mM Tris, pH 8.0) was added to the cell pellet. The pellet was briefly sonicated with a probe sonicator (QSonica, Q125 Sonicator, 125 Watt, 20 kHz) for 5 seconds and spun for a final time at 13,000 x *g* for 20 minutes at 4°C. The supernatant containing the soluble nuclear proteins was transferred to a separate labeled 1.5mL tube. The remaining pellet was discarded.

A BCA assay (Pierce™ BCA Protein Assay Kit, category number 23225) was used to determine the concentration of the resulting protein fractions. The indicated mass of protein was boiled for 5 minutes at 95°C with 4x Laemmeli Buffer (BioRad #1610747) supplemented with 2.5% (v/v) beta-mercaptoethanol and ran on a 4-20% TGX gel (BioRad #5671094 and #5671095). Protein was transferred to polyvinylidene difluoride (PVDF) membrane (BioRad #1704273) using a Trans-Blot Turbo Transfer System (BioRad). Licor Revert 700 Total protein stain (Licor, P/N: 926-11011) was used to stain the total protein on the membrane and imaged on the Odyssey FC imaging system (Licor). For the MITF and PGRN antibodies, fixation with 4% PFA in TBS for 10 minutes at RT was performed. Membranes were then blocked in 5% milk in TBS-Tw (0.1%, v/v Tween20) for an hour at room temperature (RT). Primary antibodies (PGRN R&D Systems af2557, 1: 400; GPNMB Abcam, ab188222 1:2500; MITF Cell Signaling Technologies 97800S, 1:1000; H3 Cell Signaling Technologies 9715S, 1:1000; GAPDH Cell Signaling Technologies 2118S,1:2500-5000) were diluted to the indicated concentrations in 5% milk-TBS-Tw before being applied to the membrane overnight at 4°C. Blots were washed for 5 minutes at RT using TBS-Tw three times before horseradish peroxidase (HRP)- conjugated secondary antibodies were applied for 1 hour RT. An additional three 5-minute washes removed excess secondary. Signal was elicited using SuperSignal West Femto Maximum Sensitivity Substrate (ThermoScientific™, category number 34096) and captured using Odyssey FC imaging system (Licor). ImageStudioLite was used to quantify the intensity of the bands.

The membrane was stripped between the probe of MITF and GAPDH using a buffer containing a final concentration of 1 M Tris, 20% SDS, and 0.7% BME for 15 minutes at 50°C. The stripped membrane was blocked in 5% Milk TBS-Tw for 30 minutes after stripping. An additional probe with Femto was performed to determine the success of the stripping protocol. Signal was normalized to the surrounding background before being normalized to GAPDH or H3 loading control for each band.

### RNA and qPCR

RNA was isolated using a Qiagen Mini-kit (Qiagen, category number 74104), following manufacturer instructions with minimal alterations. The concentration of BME was increased to a final concentration of 20uM to inhibit endogenous RNase activity. After elution, RNA concentration was determined using a Denovix spectrophotometer. cDNA was generated using a High-Capacity cDNA Reverse Transcription Kit (Applied Biosystems™, catalog number 4374966) following kit instructions. 2x Universal SYBR Green Fast qPCR Mix (Abclonal, RK21203) was used with the indicated primers (Table 1) for detection. A final mass of 6.25ng of cDNA was used in each well. Each sample was run in triplicate per gene. Variance above 0.4 standard deviations from the sample’s average were excluded.

**Table 1.**
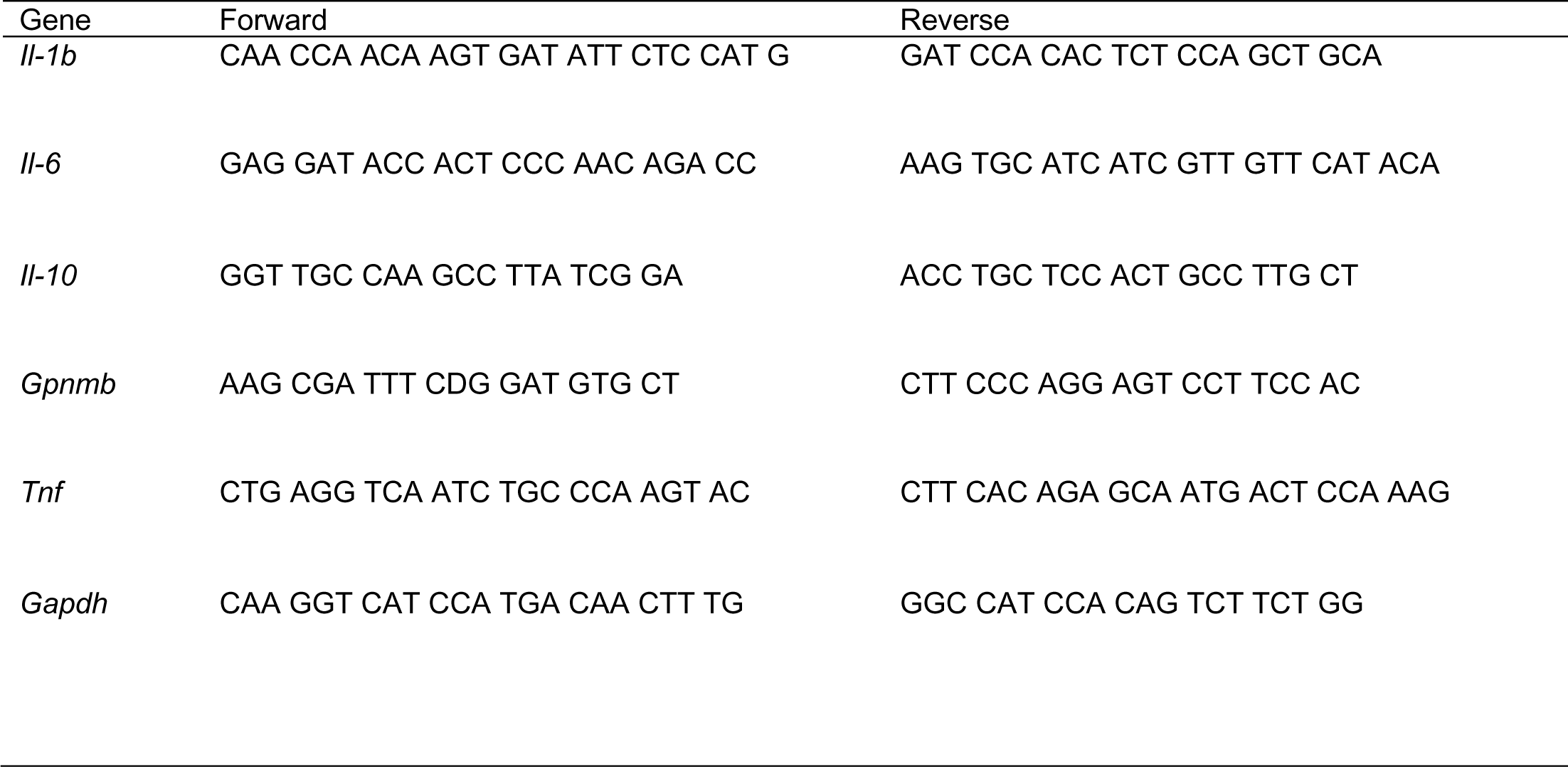
List of qPCR primers for genes of interest.

### ELISA

Conditioned media was collected from the pMacs at the indicated times. After being transferred to a 1.5mL tube, the media was centrifugated at 10,000 x *g* for 10 minutes at 4°C to remove any cell debris. The supernatant was transferred to a new tube and stored at -20°C for later analysis. Endogenous GPNMB ECF was measured using a Biotechne GPNMB ELISA Kit (Biotechne/R&D Systems, DY2330) following manufacturer instructions with minor modifications. A four-parameter logistic(4-PL) curve was generated from the GPNMB ECF standards included in the kit. The standard curve was created with an online 4-PL curve calculator from AAT Bioquest (https://www.aatbio.com/tools/four-parameter-logistic-4pl-curve-regression-online-calculator). The equation from the 4-PL curve was used to determine the concentration of GPNMB ECF in the test samples.

## Results

### GPNMB is Dysregulated in Progranulin-Deficient Macrophages

Previous work identified a novel increase in GPNMB signal in brain-resident cells, presumed to be microglia, in *Grn* KO mice at 18-months old^24^. An age series allowed the authors to conclude that there is an age-dependent increase in GPNMB signal in the central nervous system (CNS) of *Grn* KO mice. To extend on this finding, we investigated the relative abundance of GPNMB in peripheral myeloid cells by isolating thioglycollate-elicited peritoneal macrophages (pMacs) from *Grn* KO mice. Western blot analysis confirmed the absence of progranulin and also showed a significant increase in GPNMB protein in pMacs collected from 5–6-month-old *Grn* KO mice relative to B6 controls in both male and female mice (Figure 1A-C).

**Figure 1.**
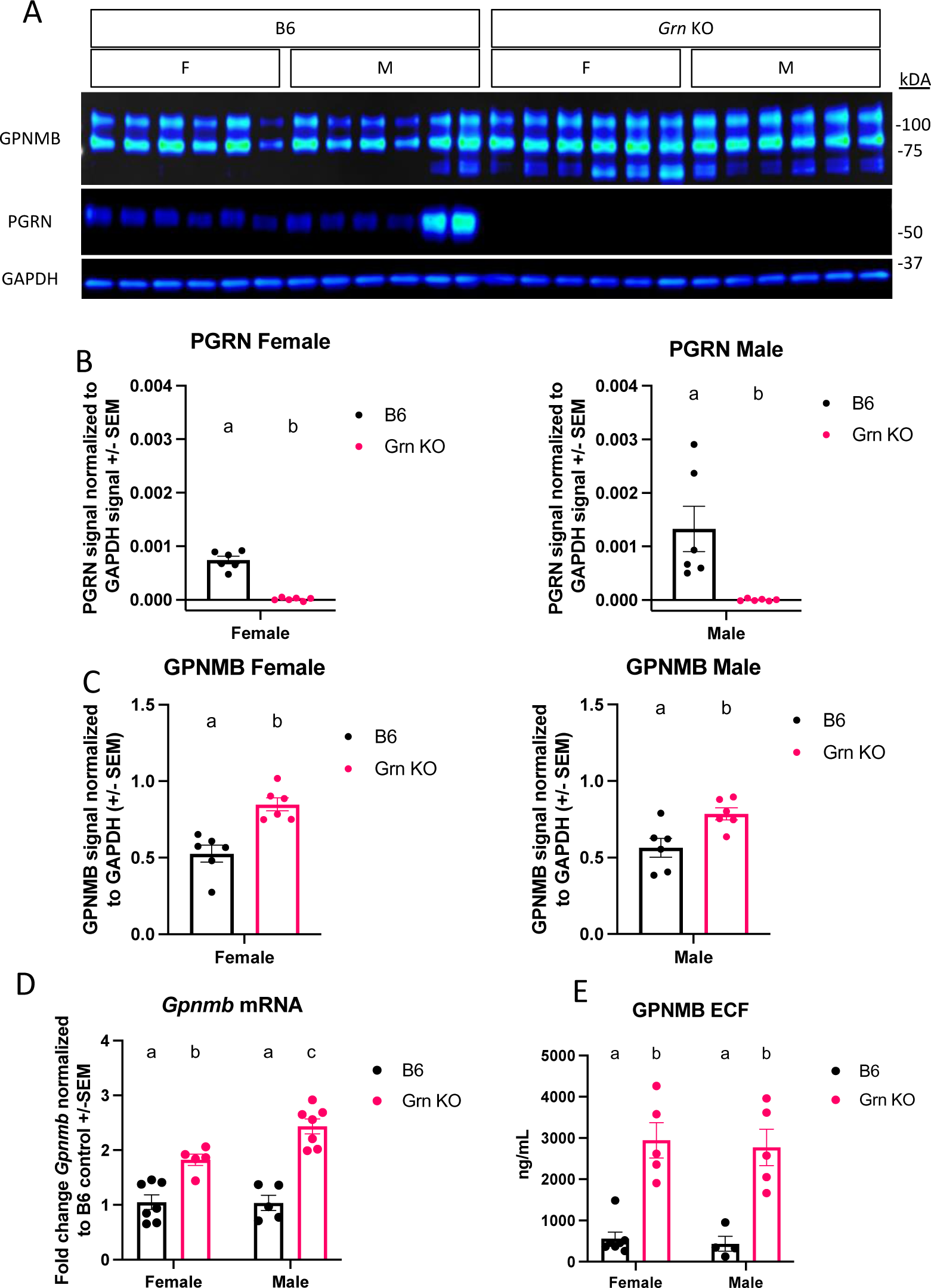
GPNMB is dysregulated in *Grn* KO pMacs. A) Representative western blot images of GPNMB, PGRN, and GAPDH from B6 and *Grn* KO pMacs B) Quantification of PGRN signal normalized to GAPDH, n=6 per sex and genotype, unpaired t-test for both females and males. C) Quantification of GPNMB signal normalized to GAPDH, n=6 per sex and genotype, unpaired t-test for both females and males. D) qPCR data on baseline *Gpnmb* transcript from both B6 and *Grn* KO pMac, n=5-7 per sex and genotype, two-way ANOVA with Sidak’s multiple comparisons test. E) GPNMB ELISA measurement of GPNMB ECF generated by pMacs during 48 hours, n=4-5 per sex and genotype, two-way ANOVA with Sidak’s multiple comparisons test. Letter(s) above the bar graphs represent the results of the *post-hoc* tests. Groups that share the same letter are not significantly different from one another.

However, it was previously unclear if the relative increase in GPNMB protein abundance was a non-specific intracellular accumulation as a result of endo-lysosomal dysfunction without sufficient progranulin expression. Using qPCR, we determined that pMacs from 5-6-month-old *Grn* KO mice had a significant increase in *Gpnmb* mRNA relative to B6 controls, with male *Grn* KO pMacs expressing significantly higher amounts than female *Grn* KO pMacs. (Figure 1D). This supports that this increase in GPNMB protein is a result of increased expression and not an endo-lysosomal bystander. Previous work has established that GPNMB is subject to cleavage events by extracellular proteases like ADAM10 that produces a soluble extracellular fragment (ECF)^26^. Given the increase in GPNMB protein substrate, we investigated the amount of GPNMB ECF that was produced in pMac media. Using an ELISA specific for endogenous GPNMB ECF, we determined that there is a significant increase in the amount of GPNMB ECF generated in pMacs collected from 5–6-month-old male and female *Grn* KO mice relative to B6 controls (Figure 1E). Together, this data demonstrates that there is an active increase in both GPNMB protein and the soluble ECF generated from it, in the absence of progranulin-deficient macrophages.

### *Grn* KO pMacs exhibit diminished expression of early inflammatory genes

Having discovered an increase in the GPNMB ECF generated by *Grn* KO pMacs, we wanted to investigate any functional effect of recombinant GPNMB extracellular domain (ECD) treatment on the behaviour of these pMacs. Previous work has identified a decrease in inflammatory gene expression with recombinant GPNMB ECD treatment in *Gpnmb* KO pMacs^28^. We adapted their paradigm to our own purposes, pre-treating two groups of cells with two doses of recombinant GPNMB ECD for four hours prior to three hours of LPS stimulation with an LPS only and no treatment control (Figure 2A). This paradigm allows us to determine the differences in inflammatory gene expression between *Grn* KO and B6 pMacs during LPS stimulation, but it also permits us to detect any changes in inflammatory gene expression with GPNMB ECD pre-treatment.

**Figure 2.**
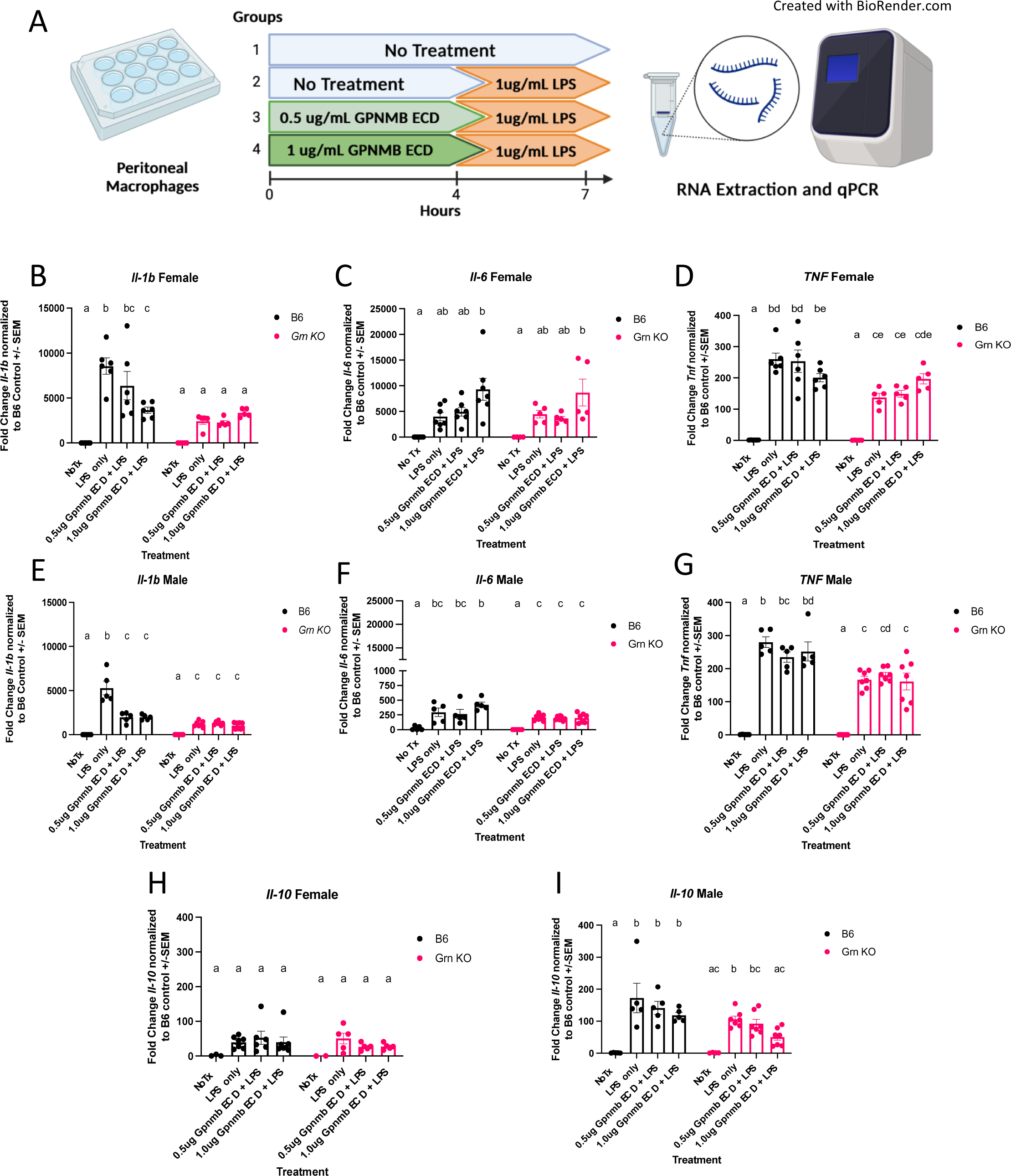
*Grn* KO pMacs show diminished expression of early inflammatory genes. A) Graphic of experimental paradigm. Created with Biorender.com. B) qPCR for *Il-1b* expression in female B6 and *Grn* KO pMacs n=5-6 per genotype C) qPCR for *Il-6* expression in female B6 and *Grn* KO pMacs, n=5-6 per genotype. D) qPCR for *Tnf* expression in female B6 and *Grn* KO pMacs, n=5-6 per genotype E) qPCR for *Il-1b* expression in male B6 and *Grn* KO pMacs, n=5-6 per genotype. F) qPCR for *Il-6* expression in male B6 and *Grn* KO pMacs, n=5-6 per genotype. G) qPCR for *Tnf* expression in male B6 and *Grn* KO pMacs, n=5-6 per genotype. H) qPCR for *Il-10* expression in female B6 and *Grn* KO pMacs, n=5-6 per genotype I) qPCR for *Il-10* expression in male B6 and *Grn* KO pMacs, n=5-6 per genotype. Two-way ANOVA with Sidak’s multiple comparisons test. Groups that share the same letter are not significantly different from one another.

qPCR analysis demonstrated multiple notable differences in the expression of inflammatory genes between genotypes. *Il-1b* transcript was increased with LPS only stimulation in both sexes of B6 pMacs, but there was a diminished if still significant increase in *Il-1b* transcript in *Grn* KO male pMacs and no change at all in *Grn* KO female pMacs with LPS only stimulation (Figure 2B and 2C). There was no significant increase in *Il-6* transcript abundance in B6 and *Grn* KO female pMacs stimulated with only LPS relative to no treatment controls, but males of both genotypes had a significant, if smaller, increase in *Il-6* transcript with only LPS stimulation (Figure 2D and 2E). *Tnf* transcript was also significantly different between the genotypes following only LPS stimulation, but neither sex significantly differed from their respective B6 controls (Figure 2F and 2G). In contrast, no differences in *Il-10* transcript between any genotype of either sex were observed (Figure 2H and 2I).

The effect of the GPNMB ECD pre-treatment is less impactful than the genotype. In both male and female B6 pMacs, there is significantly decreased *Il-1b* transcript relative to the LPS only control, suggesting that GPNMB ECD pre-treatment was sufficient to blunt the LPS-mediated increase (Figure 2B and 2C). Additionally, female pMacs from both genotypes show no significant increase in *Il-6* transcript with LPS stimulation relative to the no treatment control except for when pre-treated with 1ug/mL of GPNMB ECD (Figure 2D). However, no significant effect of GPNMB pre-treatment on *Il-6* was observed in pMacs from B6 males (Figure 2E). *Tnf* transcript levels did not significantly change with pre-exposure to either dose of GPNMB ECD (Figure 2F and 2G). Finally, *Il-10* transcript showed no change due to GPNMB ECD treatment, with the exception of the *Grn* KO males, where both doses of GPNMB ECD reduced the *Il-10* transcript to levels that are not significantly different from the no treatment control (Figure 2I).

Finally, we investigated if GPNMB ECD pre-treatment and/or LPS only treatment altered the amount of GPNMB ECF that was released into the media by pMacs. In both genotypes of female pMacs, GPNMB ECF concentration increased with LPS but did not reach statistical significance until pre-treated with 1ug of GPNMB ECD (Figure 2J). In contrast, the B6 male pMacs did not change the amount of GPNMB ECF with any treatment, but the *Grn* KO pMacs generated a significant increase in GPNMB ECF with LPS stimulation regardless of GPNMB ECD pre-treatment (Figure 2K). Collectively, this data demonstrates a key significant decrease in critical pro-inflammatory cytokine expression in *Grn* KO pMacs, and that GPNMB ECD is able to blunt the LPS-mediated increase in *Il-1b* expression in B6 pMacs

### Re-addition of progranulin rescues GPNMB phenotype in *Grn* KO pMacs

To investigate the regulation of GPNMB in progranulin-deficient macrophages, we treated both B6 and *Grn* KO pMacs with recombinant mouse progranulin. Using western blot analysis, we determined that the relative abundance of GPNMB protein was significantly increased in Grn KO pMacs that did not receive progranulin but was not significantly different from B6 controls after treatment with recombinant progranulin (Figure 3 A-D). We detected no significant change in GPNMB with the addition of exogenous progranulin in B6 pMacs (Figure 3 A-D). This data suggests that the re-addition of progranulin is sufficient to significantly decrease GPNMB levels in *Grn* KO pMacs, but the exogenous progranulin had no significant effects on the expression of GPNMB in B6 pMacs.

**Figure 3.**
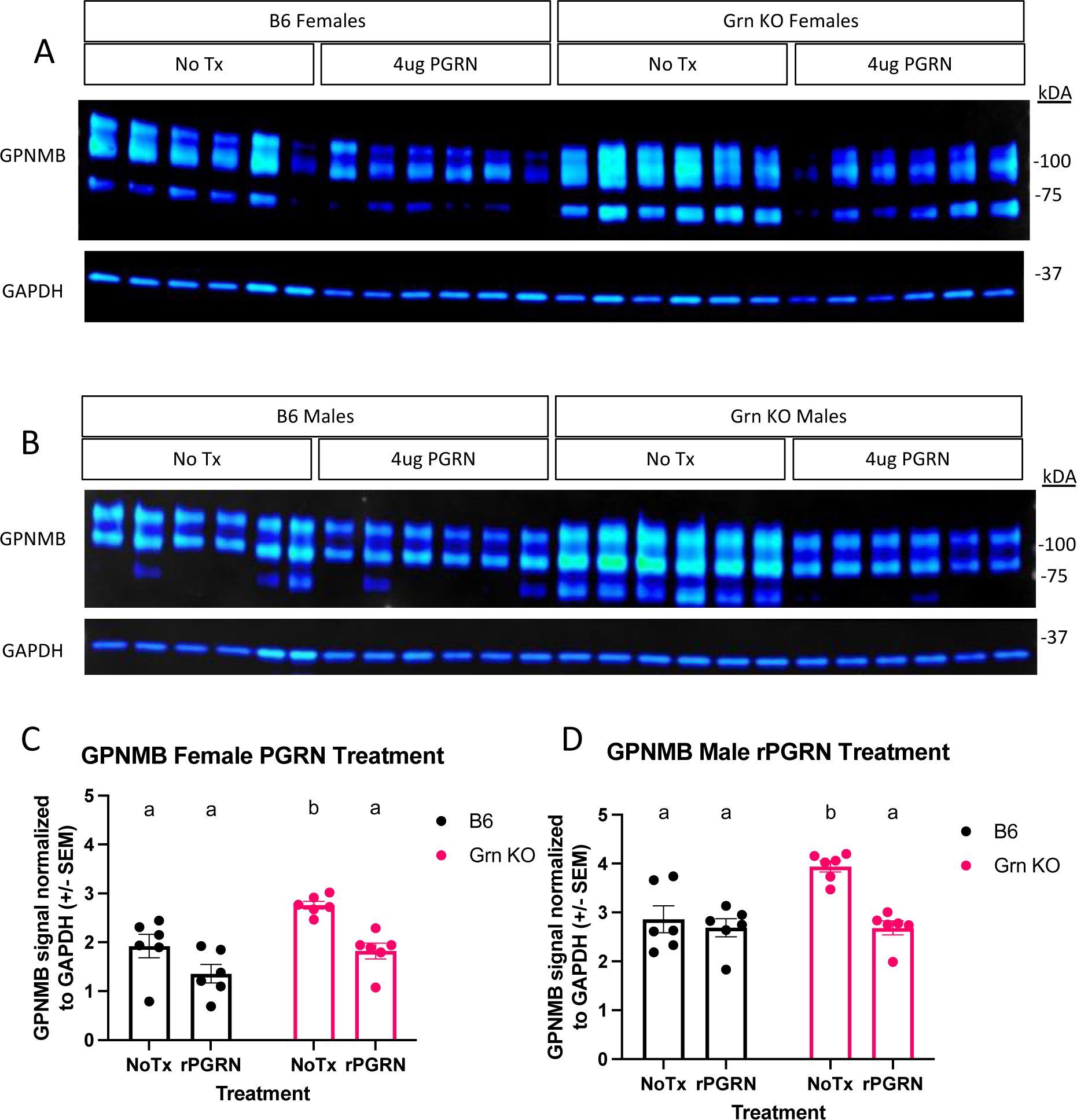
Re-addition of progranulin rescues GPNMB phenotype in *Grn* KO pMacs. A) Representative western blot images of GPNMB and GAPDH from female B6 and *Grn* KO pMacs with and without 48 hours of 4ug/mL of recombinant progranulin treatment. B) Representative western blot images of GPNMB and GAPDH from male B6 and *Grn* KO pMacs with and without 48 hours of 4ug/mL of recombinant progranulin treatment. C) Quantification of GPNMB signal normalized to GAPDH, n=6 per genotype and treatment. D) Quantification of GPNMB signal normalized to GAPDH, n=6 per genotype and treatment.

### MITF is Dysregulated in Grn KO pMacs

Previous reports have implicated the transcription factor MITF to regulate GPNMB in dendritic cells^32^, RAW264.7 cells^37^, and osteoclasts^37^. Given our observation of increased GPNMB expression in *Grn* KO pMacs, we investigated the regulation of MITF in both the presence and absence of recombinant progranulin treatment (Figure 3). To best determine the relative abundance of MITF, we used chemical fractionation to concentrate nuclear and cytosolic proteins into separate lysates. Subsequent western blot analysis identified distinct patterns of MITF abundance in the nuclear and cytosolic fractions of B6 and *Grn* KO pMacs. B6 pMacs have a greater amount of MITF in the cytosolic fraction but very little in the nuclear fraction (Figure 4 A-C). In contrast, *Grn* KO pMacs have little cytosolic MITF relative to B6, but instead have greater nuclear MITF relative to B6 pMacs. Re-addition of recombinant progranulin for 48 hours did not rescue the dysregulated nuclear MITF in *Grn* KO pMacs (Figure 4 A and C).

**Figure 4.**
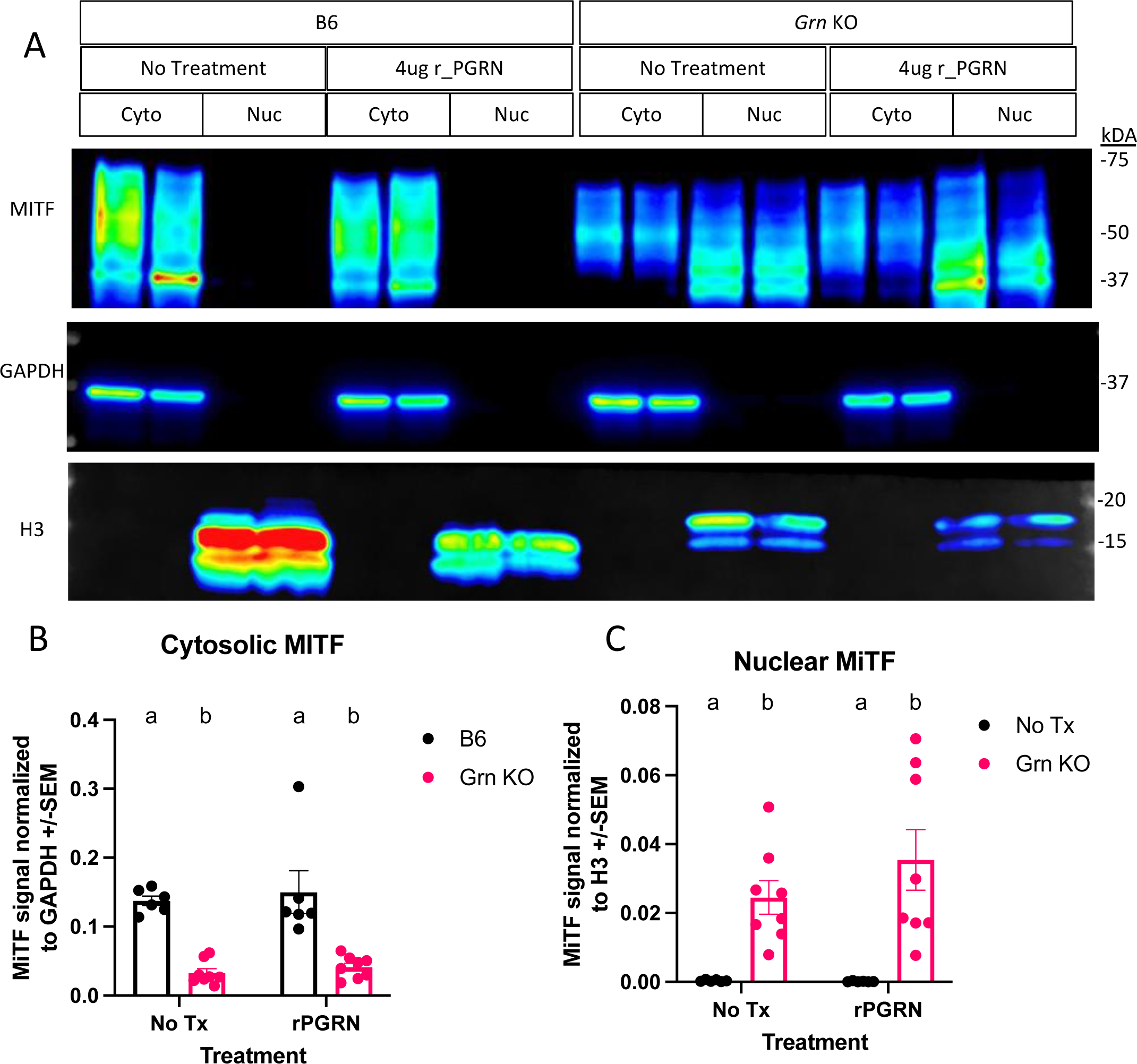
MITF is dysregulated in *Grn* KO pMacs. A) Representative western blot images of MITF, GAPDH, and H3 from B6 and *Grn* KO pMacs with and without 48 hours of 4ug/mL of recombinant progranulin treatment. B) Quantification of MITF signal normalized to GAPDH signal, n=6 per genotype, two-way ANOVA with Sidak’s multiple comparisons test. C) Quantification of MITF signal normalized to H3 signal, n=6 per genotype. Two-way ANOVA with Sidak’s multiple comparisons test. Groups that share the same letter are not significantly different from one another.

This lack of MITF rescue was initially surprising, but to better understand the underlying dynamics, we ran a western blot for progranulin itself on the cohort of pMacs treated with recombinant progranulin (Figure 5). In the 48 hours following the re-addition of progranulin, both sexes of *Grn* KO pMacs have almost completely degraded the input progranulin and show little signal beyond background in the no treatment group (Figure 5 A-D). In contrast, the female B6 pMacs that were treated with exogenous progranulin have significantly elevated cytosolic progranulin levels (Figure 5 A and C). B6 male pMacs given exogenous progranulin had no significant change from the no treatment baseline (Figure 5 B and D). The lack of MITF rescue in the *Grn* KO pMacs (Figure 3-4) correlates with the lack of available intact progranulin (Figure 5). While GPNMB was rescued with 48 hours of recombinant progranulin treatment (Figure 3), the timepoint and/or dose of progranulin was insufficient to demonstrate a progranulin-mediated rescue in MITF localization in *Grn* KO pMacs (Figures 4 and 5).

**Figure 5.**
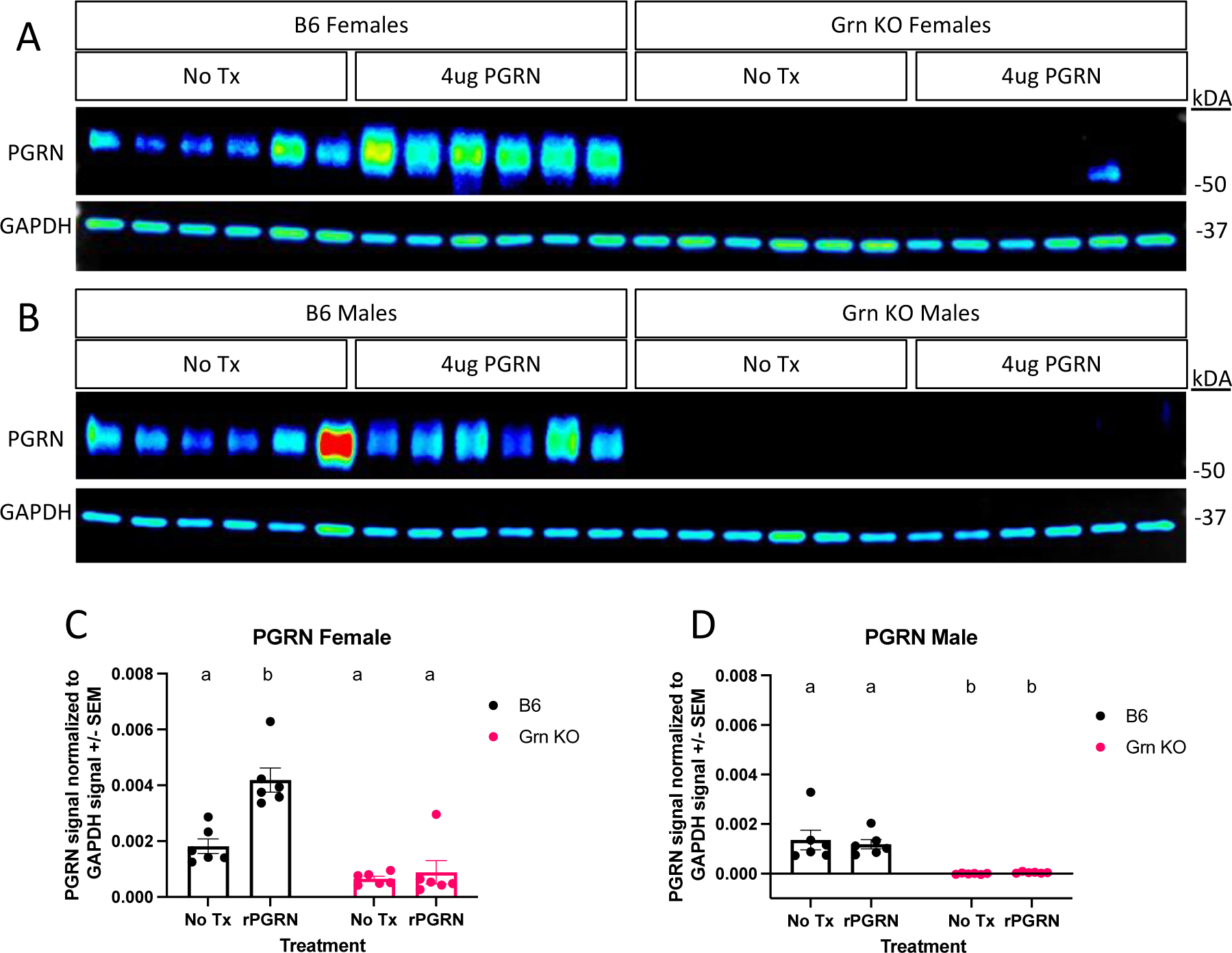
Recombinant progranulin is almost fully degraded within 48 hours by *Grn* KO pMacs. A) Representative western blot images of cytoplasmic lysate collected from female B6 and *Grn* KO pMacs in the presence or absence of 4ug/mL of recombinant mouse progranulin. B) Representative western blot images of cytoplasmic lysate collected from male B6 and *Grn* KO pMacs in the presence or absence of 4ug/mL of recombinant mouse progranulin. C) Quantification of PGRN signal normalized to GAPDH, n=6 per genotype and treatment. Two-way ANOVA with Sidak’s multiple comparisons test. Groups that share the same letter are not significantly different from one another. D) Quantification of PGRN signal normalized to GAPDH, n=6 per genotype and treatment. Two-way ANOVA with Sidak’s multiple comparisons test. Groups that share the same letter are not significantly different from one another.

## Discussion

Overall, our work demonstrates, for the first time, that GPNMB is dysregulated in peripheral macrophages in the absence of progranulin at a significantly earlier time point than previously reported in the CNS. Furthermore, we show this increase in GPNMB is driven by increased expression of *Gpnmb* and dynamically regulated by progranulin levels. Finally, we identify a novel finding of MITF dysregulation in *Grn* KO pMacs that correlates with, but does not appear to be required for, the dysregulated expression of GPNMB. Taken together, these data demonstrate the vital role of progranulin in modulating the peripheral immune environment and also highlight that events in the periphery often pre-empt those of the CNS, offering unique opportunities for early insight and intervention in the mechanism(s) of central-peripheral immune cell crosstalk that contributes to neurodegenerative disease.

First, the studies presented herein a) confirm the phenotype of increased GPNMB expression in progranulin-deficient myeloid cells previously reported^24^, and b) extend observations into the periphery revealing increases in GPNMB at a significantly earlier time point than previously reported. Additionally, our data demonstrate that this increase is a dynamic and possibly compensatory response to insufficient progranulin levels that also results in a significant increase in the concentration of the soluble signalling fragment generated from GPNMB (GPNMB ECF) which has been shown to have anti-inflammatory effects^27,28^.

Therefore, to investigate the potential effects of elevated GPNMB ECF on macrophage behaviour in response to the loss of progranulin expression, we adapted a paradigm involving pre-treatment with a recombinant fragment of the extracellular domain of GPNMB (GPNMB ECD) prior to LPS stimulation^28^. First, we found that *Grn* KO pMacs overall had reduced transcripts for classic inflammatory genes relative to B6 pMac when comparing the effect of LPS stimulation to LPS-untreated controls. This may seem counterintuitive, as progranulin is typically associated with anti-inflammatory effects, but caution is advised with interpreting an early snapshot of the inflammatory cascade as an accurate representation of the whole. A robust inflammatory response is necessary to maintain health but requires strict regulation to prevent deleterious outcomes. In our view, this lagging response to LPS stimulation in the *Grn* KO pMacs is an indication of the dysfunction related to insufficient progranulin levels. Next, pre-treatment of B6 pMacs with GPNMB ECD prior to LPS stimulation achieved a blunting of select pro-inflammatory genes like *Il-1b* in both sexes. Other early inflammatory genes did not appear to be significantly affected by the GPNMB ECD pre-treatment in B6 pMacs, but it is known that peak expression for inflammatory genes following LPS treatment or an immune challenge differ significantly^38,39^, so a single time point is insufficient to make a definitive conclusion about the possible effects that GPNMB ECD pre-treatment may have on the kinetics of inflammatory gene expression. In contrast, there was no significant effect of GPNMB ECD pre-treatment in *Grn* KO pMacs with or without LPS challenge. However, we previously showed that *Grn* KO pMacs generate significantly more GPNMB ECF than B6 pMacs while at rest so it is possible that the GPNMB ECF generated into the media between the time the cells were plated and the beginning of the stimulation paradigm affected this response and should be taken into account in the interpretation of the effects of exogenously added GPNMB ECF on inflammatory genes generated following LPS stimulation. It is possible that this inevitable “pre-pre-treatment” desensitized the *Grn* KO pMacs to any possible further response to additional GPNMB ECD administered during the LPS stimulation paradigm. Additionally, this effect may be responsible for the diminished inflammatory responses noted in *Grn* KO pMacs. Future studies will address these possibilities.

Importantly, we successfully demonstrate that progranulin re-addition is sufficient to rescue the increase in GPNMB protein in *Grn* KO pMacs, but the precise mechanism behind that rescue remains unclear and does not require normalization of MITF activity or subcellular localization. Specifically, based on previous work highlighting the role of MITF in driving the expression of CLEAR genes and the ability of MITF to drive GPNMB expression in dendritic cells, we sought to test the hypothesis that MITF was involved in the rescue mechanism. In support of our hypothesis, we found clear differences in the basal regulation of MITF in B6 versus *Grn* KO pMacs. Specifically, western blots of cytosolic and nuclear fractions revealed a clear and significant decrease in cytosolic MITF and a corresponding increase in nuclear MITF in *Grn* KO pMacs relative to B6 controls. We interpret this to represent altered MITF regulation in *Grn* KO pMacs. To further interrogate the involvement of MITF in the progranulin-mediated rescue of GPNMB expression, we then compared the baseline fractions to fractions taken from B6 and *Grn* KO pMacs treated with recombinant mouse progranulin. The same treatment was able to rescue the GPNMB phenotype in *Grn* KO pMacs, but there was no significant shift in the ratio of nuclear or cytosolic MITF in *Grn* KO pMacs that received the same treatment, suggesting that although MITF may be involved in the rescue, translocation of MITF is not required to mediate the progranulin-mediated reduction in GPNMB expression. There are several possibilities for this outcome. First, it is possible that 48 hours of 4ug/mL of progranulin treatment was not a sufficient length of time or dose of progranulin to allow significant clearance of MITF from the nucleus, and/or progranulin re-addition may have only affected the activity of MITF rather than its abundance and subcellular localization. However, there is also the possibility that MITF activity is simply not required for driving the increase in GPNMB in pMacs observed here.

Our findings represent a significant contribution to discovering additional early-stage dysregulated proteins (GPNMB and MITF) in progranulin-deficient peripheral immune cells that have known contributions to peripheral immune cell activity. GPNMB has been demonstrated to impact macrophage and T cell activity^29,30,40^ and while the mechanism(s) are unclear, GPNMB has been previously associated with risk for PD ^18,41^ and, with some controversy, found to be increased in AD patients and mouse models of AD^42-46^. MITF also has significant association to immune cells. Multiple co-factors and transcription factors, including PU.1, and NFATc1, interact with MITF, influencing the differentiation of immune cells^47-49^. Elucidating the consequences of MITF dysregulation on the composition and integrity of progranulin-deficient peripheral immune cell populations is a novel avenue of research that will permit further insight into the role of central-peripheral crosstalk in neurodegenerative diseases.

## Acknowledgements

We thank the Tansey lab for their feedback concerning this review.

## Author contributions

DAG, RLW, and MGT designed the experiments. DAG, RLW, and MGT wrote and edited the manuscript. NKN and CC performed husbandry and genotyping. DAG created the figures. All authors read and approved the final manuscript.

## Funding

The authors are supported by Grant RF1NS28800 from the National Institute of Health and the National Institute of Neurological Disorders and Stroke (MGT) and an Alzheimer’s disease and related dementias (ADRD) teaching fellowship funded by the National Institute of Health (T32AG061892) at the University of Florida (DAG).

## Disclosures

The authors declare no conflicts.

## Notes

### Competing Interest Statement

The authors have declared no competing interest.

